# An obligatory role for AgRP neurons in maintaining body temperature during time-restricted feeding

**DOI:** 10.64898/2025.12.16.694714

**Authors:** Cunjin Su, Jing Cai, Yuanzhong Xu, Benjamin R. Arenkiel, Qingchun Tong

## Abstract

Homeotherms maintain a steady body temperature through thermoregulation, a process critical for survival during fasting, in which the brain has to defend energy-costly body temperature while reducing energy expenditure to conserve energy reserve; however, the neural basis for defending body temperature remains unclear. Here, we demonstrated that AgRP neuron lesion led to lethality during time-restricted feeding but no obvious impact on HFD-induced obesity or obesity-reducing responses to glucagon-like peptide-1 receptor agonism. The lesion disrupted adaptive feeding behaviors during time-restricted feeding and reduced motivational feeding. Notably, the lethality was caused by hypothermia instead of reduced food intake. The lesion also caused failure in body temperature maintenance during acute fasting in cold. Fasting-induced activation in AgRP neurons was abrogated when mice were placed in a warm environment. Our results identify the physiological role for AgRP neurons in defending body temperature during restricted feeding but dispensable for body weight regulation with food *ad libitum*.

## Introduction

Homeotherms maintain their body temperature within a steady range, which is higher than the ambient temperature, and as a consequence requires a significant energy cost, even at rest (Reitman, 2018). The maintenance of body temperature is critical for survival, especially when animals face a challenging environment including fasting and cold exposure (Reitman, 2018). Therefore, rigorous brain mechanisms exist to defend body temperature from deviating from the normal range. In this regard, fasting are known to increase hunger to seek energy intake and reduce metabolism to conserve energy for survival (Cannon and Nedergaard, 2004; Deem et al., 2021; Reitman, 2018). Along this line, previous studies have shown that fasting induces robust activation of brain neurons including arcuate neurons in the hypothalamus that function to promote hunger and reduce energy expenditure (Douglass et al., 2023; Jensen et al., 2013). Likewise, cold exposure activates a neural network, mainly those in the preoptic area, to increase heat production through thermogenesis and prevent heat loss through vasoconstriction (Tabarean et al., 2010; Tan and Knight, 2018). Despite intensive studies on physiological responses to cold exposure and fasting, how the brain coordinates feeding and thermogenesis to maintain body temperature balance during chronic restricted feeding regimens (Gallop et al., 2023; Wheatley et al., 2025), procedures widely reported to benefit metabolic health, remains unclear.

Arcuate neurons co-expressing agouti-related protein (AgRP) and neuropeptide Y, one of the key component neuron groups in the melanocortin pathway, are known to be a key regulator in feeding and related adaptive behaviors to environmental changes (Deem *et al*., 2021). It is well-established that AgRP neurons are sufficient to drive feeding and obesity development as evidenced by rapid feeding behaviors elicited by optogenetic and chemogenetic activation of AgRP neurons (Aponte et al., 2011; Krashes et al., 2011), and development of massive obesity by chronic activation of these neurons through targeted deletion of leptin receptors or expression of neuron-activating channels (Xu et al., 2018; Zhu et al., 2020). Thus, AgRP neurons, when activated, are poised to promote positive energy balance. Consistent with this, fasting is known to induce robust activation of AgRP neurons, and mice with AgRP neuron lesion exhibit reduced fasting-induced hyperphagia (Cai et al., 2023), suggesting an importance of AgRP neuron activation in response to fasting in behavioral adaption to environmental challenges with food insufficiency. Aside from responses to fasting, AgRP neurons are able to sense ambient temperature and regulate feeding (Deem et al., 2020). Consistently, AgRP neurons receive projections from neurons in the preoptic area, a brain region known to regulate body temperature and sense changes in environmental temperature (Yang et al., 2021). However, in addition to feeding regulation, whether and how AgRP neurons modulate body temperature is unknown.

Contrary to the previous belief that the presence of AgRP neurons is essential for survival even with *ad libitum* feeding conditions (Luquet et al., 2005), our recent study with nearly complete lesion of AgRP neurons in the brain of adult mice surprisingly caused no impact on *ad libitum* feeding or body weight (Cai *et al*., 2023). Notably, this observation is supported by other studies with chronic or chemogenetic inhibition of AgRP neurons showing no or minimal impact on body weight (Uner et al., 2019; Zhu *et al*., 2020), or with complete neurotransmission blockage of these neurons having no impact on body weight or obesity in leptin deficient *ob/ob* mice (Cavalcanti de Albuquerque et al., 2025; Gou et al., 2025). Along this line, AgRP neurons have been suggested to be required for normal body weight regulation in various nutrient-challenging conditions including high-fat high caloric diet (HFD) feeding, albeit still with *ad libitum* feeding conditions (Cai *et al*., 2023; Miletta et al., 2020; Padilla et al., 2016; Tan et al., 2014). Specifically, previous studies suggested that mice with gene manipulations in AgRP neurons reduce feeding in HFD-induced obesity (DIO) (Ajwani et al., 2024; Quiñones et al., 2018; Üner et al., 2015; Zhang et al., 2024). These results collectively suggest that specific manipulations of AgRP neurons, presumably those leading to inhibition of AgRP neuron activity, are able to confer resistance to DIO. In addition, AgRP neurons have also been suggested to function as downstream neurons that mediate obesity-reducing effects of glucagon-like peptide-1 receptor agonism (GLP-1RA) (Kim et al., 2024; Webster et al., 2024), therefore implicating a role for AgRP neurons in resisting DIO and mediating GLP-1R based obesity therapies. On the other hand, loss of AgRP neurons or deletion of NPY from AgRP neurons has been reported to paradoxically increase body weight (Denis et al., 2015; Joly-Amado et al., 2012; Qi et al., 2022). Thus, the role of AgRP neurons in *ad libitum* feeding conditions, especially when fed HFD, requires critical further clarification.

To determine whether AgRP neurons are required for adaptive responses in both feeding and temperature in time-restricted feeding, a condition mimicking food-insufficient environments, and body weight regulation in HFD *ad libitum* conditions mimicking over-nutrition environments, we employed mice with nearly complete lesion of AgRP neurons in the adult age (Cai *et al*., 2023). These mice in principle represents maximal loss of function of these neurons without a concern for potential functional compensation during development. Our results demonstrate that these mice succumbed to death within several days when challenged with time-restricted feeding while exhibited no difference in changes of body weight when challenged with HFD or GLP1-RAs, highlighting a selective requirement of AgRP neurons in adaptive feeding behaviors under food-insufficient environments. Mechanistically, lesions of AgRP neurons reduced motivational feeding and caused less feeding during restrict feeding periods. Importantly, the lethality was shown to be caused by hypothermia instead of less feeding. In addition, mice with AgRP neuron lesion exhibited failure in body temperature maintenance during cold exposure in fasting. When mice were placed in a warm environment, fasting-induced hyperphagia was blunted and fasting-induced activation of AgRP neurons was completely abrogated, suggesting a physiological role for AgRP neurons in promoting basal metabolic rate for body temperature maintenance by increasing energy intake. This line of evidence collectively demonstrates an obligatory role for AgRP neurons in defending body temperature maintenance during chronic feeding restriction.

## Results

### Nearly complete AgRP neuron lesions in adult mice caused no alterations in response to DIO or GLP-1RA

Given the strong existing literature supporting a role for AgRP neurons, when with reduced neuron activity, in rendering resistance to DIO (Ajwani *et al*., 2024; Quiñones *et al*., 2018; Üner *et al*., 2015; Zhang *et al*., 2024), we aimed to examine body weight responses in mice with AgRP neuron lesion in the adult age on HFD feeding. We used our previously established protocol to achieve effective lesion of AgRP neurons in adult mice by icv injection of 5ng DT (Cai *et al*., 2023). Consistent with the previous result, the DT administration led to nearly complete lesion of AgRP/NPY neurons (Figs. 1A and Supplementary Fig. 1). When fed HFD, these mice were monitored for weekly body weight, which was not different from littermate controls within the 10-week monitoring period in males (Fig. 1B) or females (Fig. 1D). In line with these results, no differences were found in food intake (Figs. 1C and 1E), O2 consumption (Supplementary Figs. 2A and 2D), or locomotor activity (Supplementary Figs. 2C, 2B, 2E, 2F) between the groups when fed either chow (Supplementary Figs. 2A-2C) or HFD (Supplementary Figs. 2D-2F). Importantly, feeding amounts and patterns were also comparable between groups when fed either chow (Supplementary Fig. 2G) or HFD (Supplementary Fig. 2H). These data suggest that AgRP neurons are not required in normal energy balance responses to HFD feeding.

**Figure 1.**
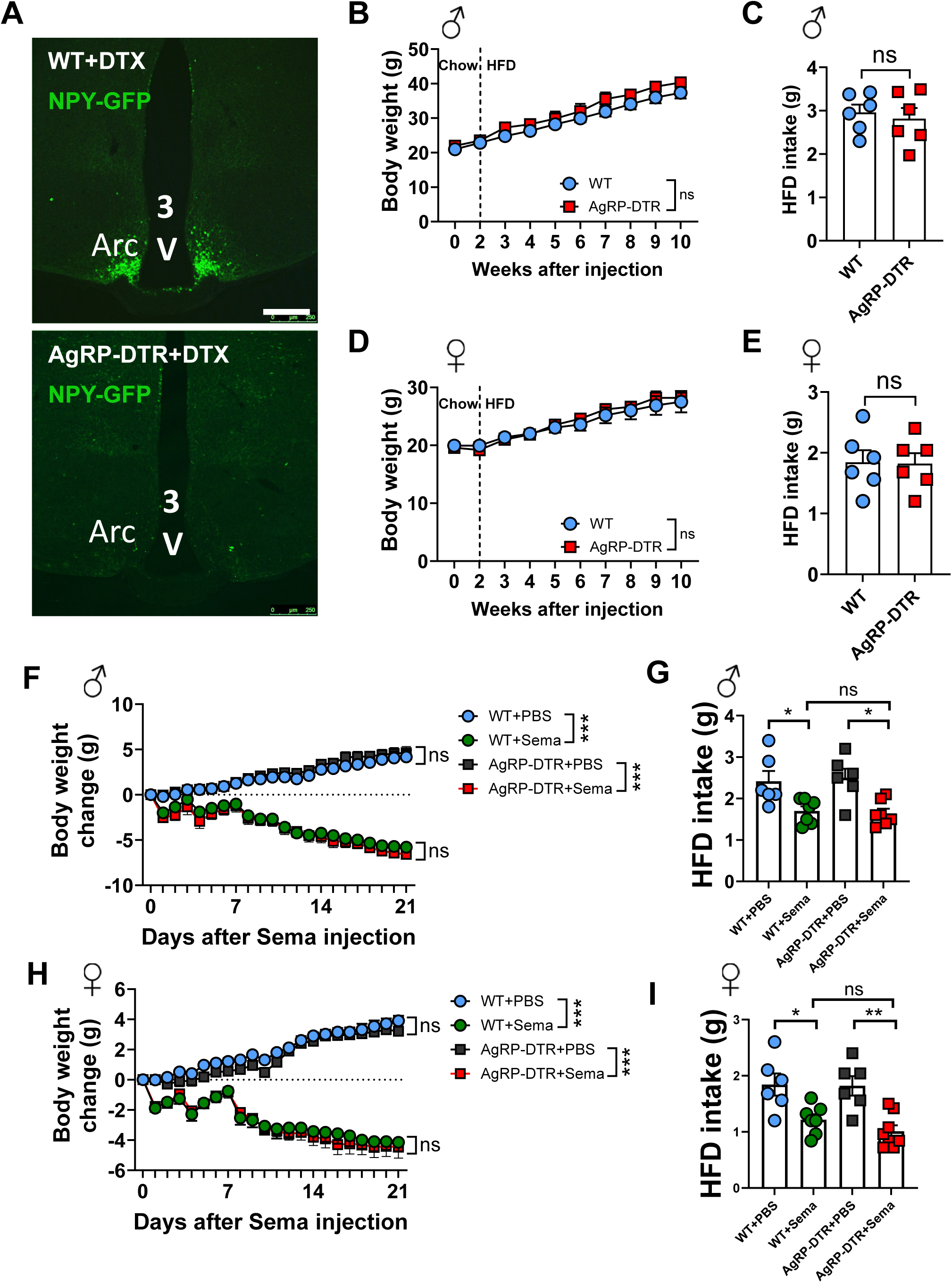
Near complete AgRP neuron lesions in adult mice caused no alterations in response to DIO and GLP1-RAs in reducing obesity. **(A)** Representative pictures showing NPY-GFP (green) positive neurons in the arcuate of WT (NPY-GFP) and AgRP-DTR (AgRP-DTR::NPY-GFP) mice that received a single i.c.v injection of 5 ng DTX. Scale bar: 250 μm. **(B)** Male mice were put on HFD two weeks after DTX injection, body weight was recorded weekly in the following 8 weeks. Two-way repeated ANOVA followed by Bonferroni’s multiple comparisons; p>0.05 between two groups. n=8 for WT group, n=5 for AgRP-DTR group. **(C)** Daily HFD intake of WT and AgRP-DTR male mice. Unpaired t-test, p=0.645 between two groups. n=6 per group. **(D)** Female mice were put on HFD two weeks after DTX injection, body weight was recorded weekly in the following 8 weeks. Two-way repeated ANOVA followed by Bonferroni’s multiple comparisons; p>0.05 between two groups. n=8 for WT group, n=15 for AgRP-DTR group. **(E)** Daily HFD intake in WT and AgRP-DTR female mice. Unpaired t-test, p=0.931. n=6 per group. Two weeks after a single i.c.v injection of 5 ng DTX, male and female WT and AgRP-DTR mice were put on HFD for 10 weeks, then they were received twice subcutaneous (s.c) injection of 40 µg/kg semaglutide for one week and daily for the next two weeks. **(F)** Body weight of male WT and AgRP-DTR mice was measured daily during semaglutide treatments. Two-way repeated ANOVA followed by Bonferroni’s multiple comparisons test, p>0.05 between WT-semaglutide and AgRP-DTR-semaglutide group. ***p<0.001 between the indicated groups. n=6 for WT+PBS and AgRP-DTR+PBS groups, n=7 for WT+Sema and AgRP-DTR+Sema groups. **(G)** Daily HFD intake of male WT and AgRP-DTR mice was measured during semaglutide treatments. Two-way repeated ANOVA followed by Bonferroni’s multiple comparisons test, p=0.049 between WT-PBS group and WT-semaglutide group, p= 0.013 between AgRP-DTR-PBS group and AgRP-DTR-semaglutide group. n=6 for WT+PBS and AgRP-DTR+PBS groups, n=7 for WT+Sema and AgRP-DTR+Sema groups. **(H)** Body weight of female WT and AgRP-DTR mice was measured daily during semaglutide treatment. Two-way repeated ANOVA followed by Bonferroni’s multiple comparisons test, p>0.05 between WT-semaglutide and AgRP-DTR-semaglutide group. ***p<0.001 between the indicated groups. n=6 for WT+PBS and AgRP-DTR+PBS groups, n=7 for WT+Sema group, n=8 for AgRP-DTR+Sema group. **(I)** Daily HFD intake of female WT and AgRP-DTR mice was measured during semaglutide treatment. Two-way repeated ANOVA followed by Bonferroni’s multiple comparisons test. p=0.033 between WT-PBS group and WT-semaglutide group, p=0.003 between AgRP-DTR-PBS group and AgRP-DTR-semaglutide group. n=6 for WT+PBS and AgRP-DTR+PBS groups, n=7 for WT+Sema group, n=8 for AgRP-DTR+Sema group. ns, not significant. All data represent the mean ± SEM.

The arcuate nucleus has been shown to be the prime site with GLP1-RA binding once administered systematically and has been implicated in responding to GLP1-RAs (Dong et al., 2021; Kim *et al*., 2024; Secher et al., 2014). Along this line, AgRP neurons, as one major group of the acuate nucleus, respond to GLP1-RAs in neuron activity (Dong *et al*., 2021). However, whether these neurons are required for the anti-obesity effect of GLP1-RAs remains untested. To specifically examine this, we used obese AgRP-DTR and littermate control male mice that have been fed HFD for 10 weeks and subjected them with Semaglutide treatments. Both groups exhibited body weight reduction indistinguishably (Fig. 1F) with a comparable degree of reduction in HFD food intake (Fig. 1G). Similar results were also observed in females in response to Semaglutide including body weight reduction (Fig. 1H) and HFD feeding (Fig. 1I). These results demonstrate that arcuate AgRP neurons are not required for Semaglutide in reducing DIO.

### AgRP neuron lesion is incompatible for survival in chronic food restriction

Lesions of AgRP neurons blunt fast-induced hyperphagia (Cai *et al*., 2023), suggesting a role for these neurons in mediating adaptive behaviors during food-insufficiency. To examine whether AgRP neurons are required for adaption to long-term food insufficiency, we subjected a group of chow-fed AgRP-DTR and control mice to a food restriction regimen, in which chow food was made only available during the night time (Fig. 2A). Interestingly, compared to control males, AgRP-DTR male mice exhibited a lethal phenotype with all mice succumbed to death during the first 8 days with the restricted feeding schedule (Fig. 2B), which was associated a dramatic reduction in body weight (Fig. 2C). Interestingly, while control mice gradually increased their food intake during the feeding period, AgRP-DTR mice exhibited a continuing reduction in food intake (Fig. 2D), suggesting an inability in adapting to food restricted environments.

**Figure 2.**
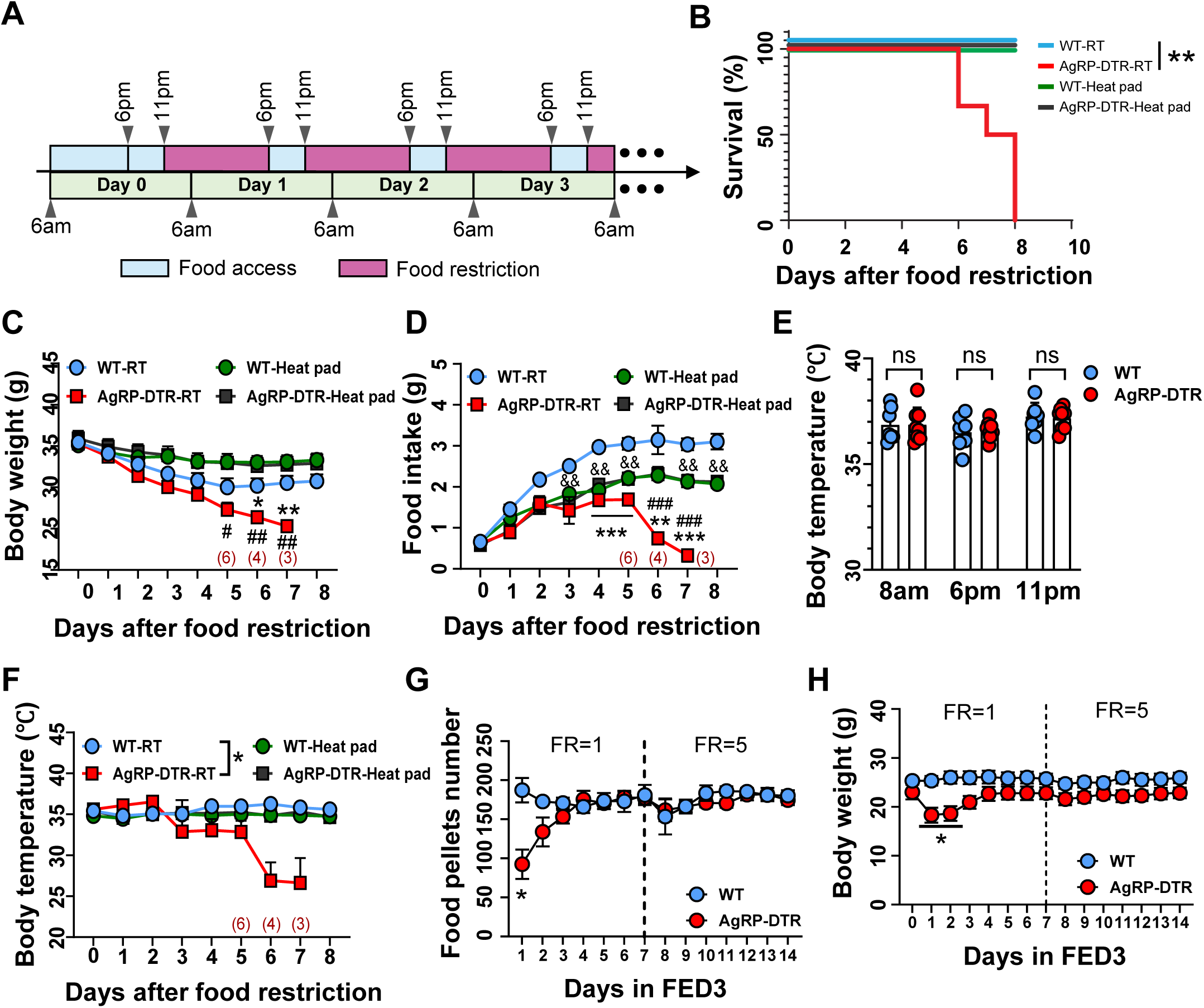
AgRP neuron lesion is incompatible for survival in chronic food restriction. Three weeks after a single i.c.v injection of 5 ng DTX, WT and AgRP-DTR male and female mice were subjected to a food restriction, whereby the mice were granted access to food only during 6 pm to 11 pm. **(A)** Schematic of food restriction paradigm. **(B-E)** Survival curve of mice (B), body weight (C), food intake (D), body temperature (E) of male WT and AgRP-DTR mice after a single i.c.v injection of 5 ng DTX was recorded during the food restriction treatment. (B) Log-rank test, p=0.002 between WT-RT and AgRP-DTR-RT groups. (C, D) Two-way repeated ANOVA followed by Tukey’s multiple comparisons test, *p<0.05, **p<0.01, ***p<0.001 between AgRP-DTR-RT and WT-RT groups; #p<0.05, ##p<0.01, ###p<0.001 between AgRP-DTR-RT and AgRP-DTR-Heat pad groups; &&p<0.01 between WT-RT and WT-Heat pad groups. (E) Two-way repeated ANOVA followed by Bonferroni’s multiple comparisons test, p<0.05 between WT-RT and AgRP-DTR groups. n=5 for WT-RT group, n=6 for AgRP-DTR-RT / WT-Heat pad / AgRP-DTR-Heat pad groups. Red numbers indicate the number of surviving mice at each corresponding time point. **(F)** Body temperature of WT and AgRP-DTR mice measured at different time points under ad libitum feeding before food restriction at room temperature. ns, not significant. **(G, H)** WT and AgRP-DTR were challenged with FED3. We used a fixed-ratio of 1 (FR=1, 1 nose poke for 1 food pellet) for the first week and of 5 (FR=5, 5 nose poke for 1 food pellet) for the second week; and comparisons were shown for feeding (G) and body weight (H). Two-way repeated ANOVA followed by Bonferroni’s multiple comparisons test, *p<0.05. n=6 for WT group, n=7 for AgRP-DTR group. All data represent the mean ± SEM.

Previous studies suggest that AgRP neurons are responsive to temperature changes (Deem *et al*., 2020). We therefore also monitored body temperature on these mice and found that there was no change in body temperature throughout 3 time points measured during a day with food provided *ad libitum* (Fig. 2E). Interestingly, when body temperature was measured during the food restriction study, we found that the lesion mice exhibited a dramatic reduction in body temperature starting day 6 after the food restriction (Fig. 2F), which coincides with the time when these mice showed reduced food intake (Fig. 2D). To selectively examine the role of the observed hypothermia, we added two additional groups of control and lesion mice with the exception that these mice were maintained on a heating pad (36°C) to prevent body temperature drop. The heating pad treatment effectively rescued the lethality (Fig. 2B), hypophagia (Fig. 2D) and hypothermia (Fig. 2F). Consistent with the rescuing effects on food intake and survival, the heating pad treatment rescued body weight drop (Fig. 2C). There results collectively suggest that AgRP neurons are required for body temperature maintenance in a food-insufficient environment.

To further probe the observed inability of AgRP-DTR mice in adapting to eat more food during the food availability period, we subjected these mice to the FED3 feeding apparatus, which requires mice to use nose pokes to obtain food pellets. We used a fixed ration of 1 (i.e. 1 nose poke for 1 food pellet) for the first week and of 5 for the second week. AgRP-DTR mice consumed significantly less food pellets during the initial 2 days of testing and gradually recovered to the control level (Fig. 2G). Consistently, their body weight was reduced during the same period (Fig. 2H). These results suggest that AgRP neuron lesion causes an acute defect in motivational feeding, which may underlie reduced feeding during feeding restriction.

### Lethality in restricted feeding is due to hypothermia

Given the association of the observed lethality with both hypophagia and hypothermia in mice with AgRP lesion, we next examined the effect of hypothermia by a pair-feeding study. We used the same restricted feeding paradigm but only provided the amount of food consumed in AgRP-DTR mice housed in room temperature (AgRP-DRT-RT) to AgRP-DTR mice paced on a heating pad (AgRP-DTR-pad) and wild type controls housed in room temperature (WT-RT) (Fig. 3A). All 3 groups consumed the same amount of food intake during the duration of the study (Fig. 3B). As expected, AgRP-DTR-RT mice died within 10 days after the start of the restricted feeding protocol. Interestingly, the pair-fed AgRP-DTR-pad mice survived similarly to the WT-RT group (Fig. 3C). Consistently, in contrast to a rapid drop in body weight (Fig. 3D) and body temperature (Fig. 3E) in AgRP-DTR-RT mice, AgRP-DTR-pad mice maintained their body weight and body temperature (Figs. 3D and 3E). There results suggest that the lethality caused by AgRP neuron lesion in restricted feeding is due to hypothermia instead of reduced food intake. Given the same amount of food consumption between AgRP-DTR-RT and AgRP-DTR-pad groups, these results also suggest that AgRP neurons are required for feeding-induced body temperature maintenance.

**Figure 3.**
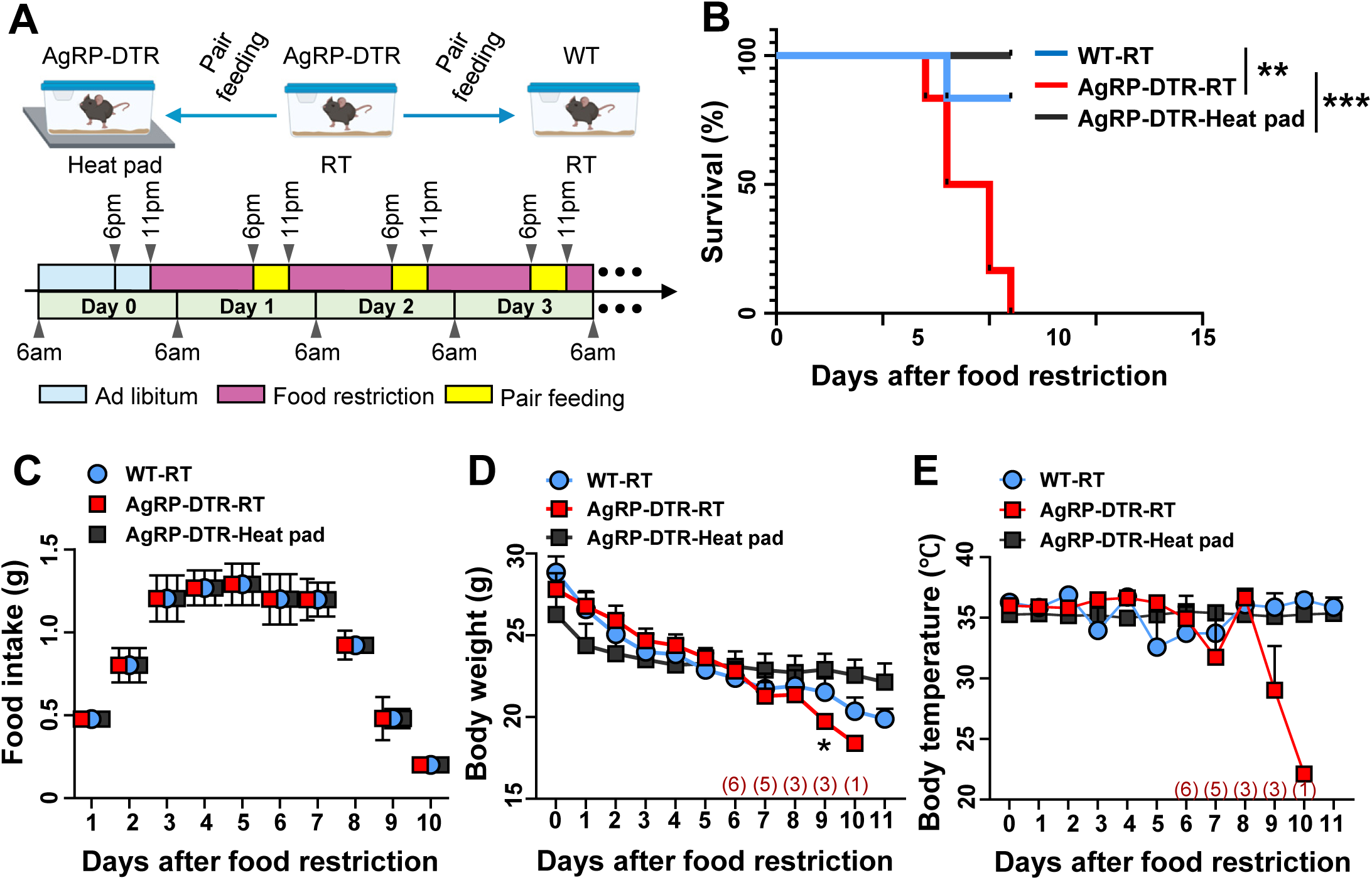
Lethality in restricted feeding is due to hypothermia. **(A)**Schematic of food restriction paradigm. The mice in WT-RT and AgRP-DTR-Heat pad groups were pair-fed to mice in AgRP-RT group. **(B)** Survival curve during pair feeding. Log-rank test, p=0.006 between AgRP-DTR-RT and WT-RT groups; p<0.001 between AgRP-DTR-RT and AgRP-DTR-Heat pad groups. n=6 for each group. **(C-E)** Food intake (C), body weight (D), and body temperature (E) during pair feeding. (D) Two-way repeated ANOVA followed by Tukey’s multiple comparisons test, *p<0.05 between AgRP-DTR-RT and AgRP-DTR-Heat pad groups. n=6 for each group. Red numbers indicate the number of surviving mice at each corresponding time point. All data represent the mean ± SEM.

### AgRP neurons are required for body temperature maintenance in cold

Given our existing data on the effect of chronic restricted feeding in body temperature with AgRP neuron lesion and previous results on the involvement of AgRP neurons in normal response to cold(Deem *et al*., 2020), we further examined feeding and body temperature responses in acute cold exposure in mice with AgRP neuron lesion. We subjected mice with AgRP neuron lesion and controls to a 4°C degree challenge for 5 hours with fasting or food provided ad libitum (Fig. 4A). Consistently with previous results (Cai *et al*., 2023), there was no difference in food intake when mice were housed in room temperature (Fig. 4B). In contrast, compared to controls, food intake was significantly reduced in mice with AgRP neuron lesion when mice were housed in cold (Fig. 4B), suggesting that AgRP neurons are required for cold-induced hyperphagia. Interestingly, responses in body temperature in these mice were not altered compared to control mice in either housing conditions (Fig. 4C), suggesting that AgRP neurons are not required for body temperature maintenance in acute exposure to cold when food provided *ad libitum*. In contrast, during the same cold exposure, when food was not provided, the lesioned mice exhibited a significant reduction in body temperature compared to controls (Fig. 4D). These results collectively suggest that food intake in these mice, even at a reduced level, is both required and sufficient for AgRP neurons to mount a normal body temperature response to cold.

**Figure 4.**
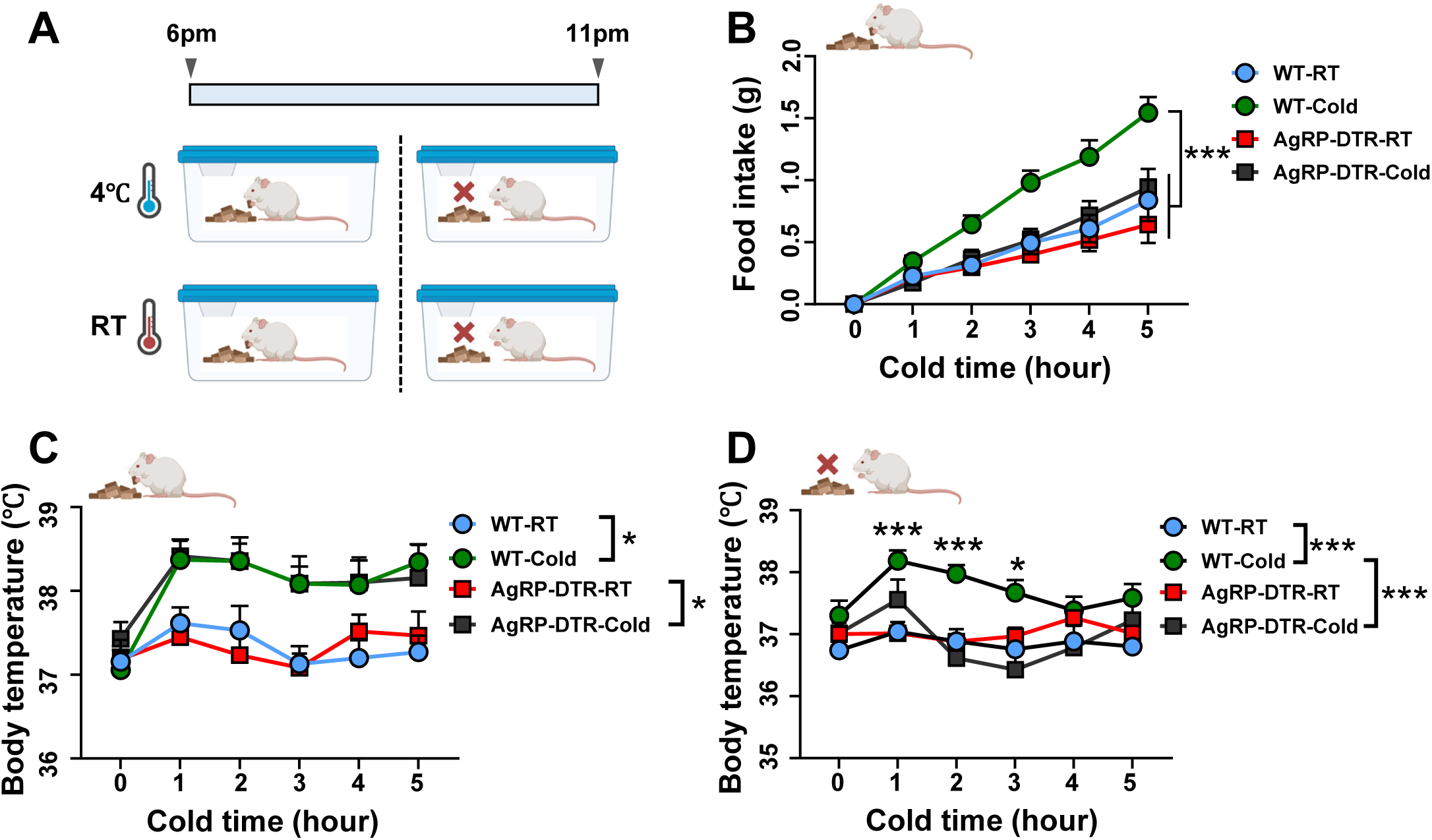
AgRP neurons are required for body temperature maintenance in cold. **(A)** Schematic of acute cold exposure paradigm. **(B)** Food intake and **(C)** body temperature of WT and AgRP-DTR mice were measured during exposure to 4 °C from 6 PM to 11 PM in the presence of food. Two-way repeated ANOVA followed by Bonferroni’s multiple comparisons test, (B) p<0.001 between WT-cold and each of the other three groups, (C) p<0.001 between WT-RT and WT-cold groups; p<0.001 between AgRP-DTR-RT and AgRP-DTR-cold groups. **(D)** body temperature of WT and AgRP-DTR mice were measured during exposure to 4 °C from 6 PM to 11 PM without food. Two-way repeated ANOVA followed by Bonferroni’s multiple comparisons test, p<0.001 between WT-RT and WT-cold groups, p<0.001 between WT-cold and AgRP-DTR-cold groups. n=7 for WT-RT/ WT-cold/ AgRP-DTR-cold groups, n=6 for AgRP-DTR-RT group. All data represent the mean ± SEM.

### Contribution of AgRP neurons in energy intake to body temperature maintenance during fasting

To further ascertain the relationship between AgRP neurons and body temperature, we test fasting-induced feeding in AgRP-DTR and control mice when mice were placed in room temperature conditions or on a heating pad (Fig. 5A). As expected, AgRP-DTR mice exhibited reduced fasting-refeeding when housed at room temperature (Fig. 5B). However, with heating pad, they showed no difference in fasting-refeeding (Fig. 5B), which is mainly due to reduced fasting-refeeding in controls mice with heating pad, presumably due to less basal metabolic rate being required to maintain a difference between body temperature and room temperature. To test this possibility, we placed mice at room temperature during fasting period and switched to heating pad only during the feeding testing period (Fig. 5C), which would eliminate the difference from body temperature maintenance during the fasting period. During this condition, no difference in fasting-refeeding was observed between the groups in either room temperature or heating pad conditions, confirming that the reduction in food intake control mice in heating pad conditions relative to room temperature is due to less energy expenditure required for body temperature maintenance.

**Figure 5.**
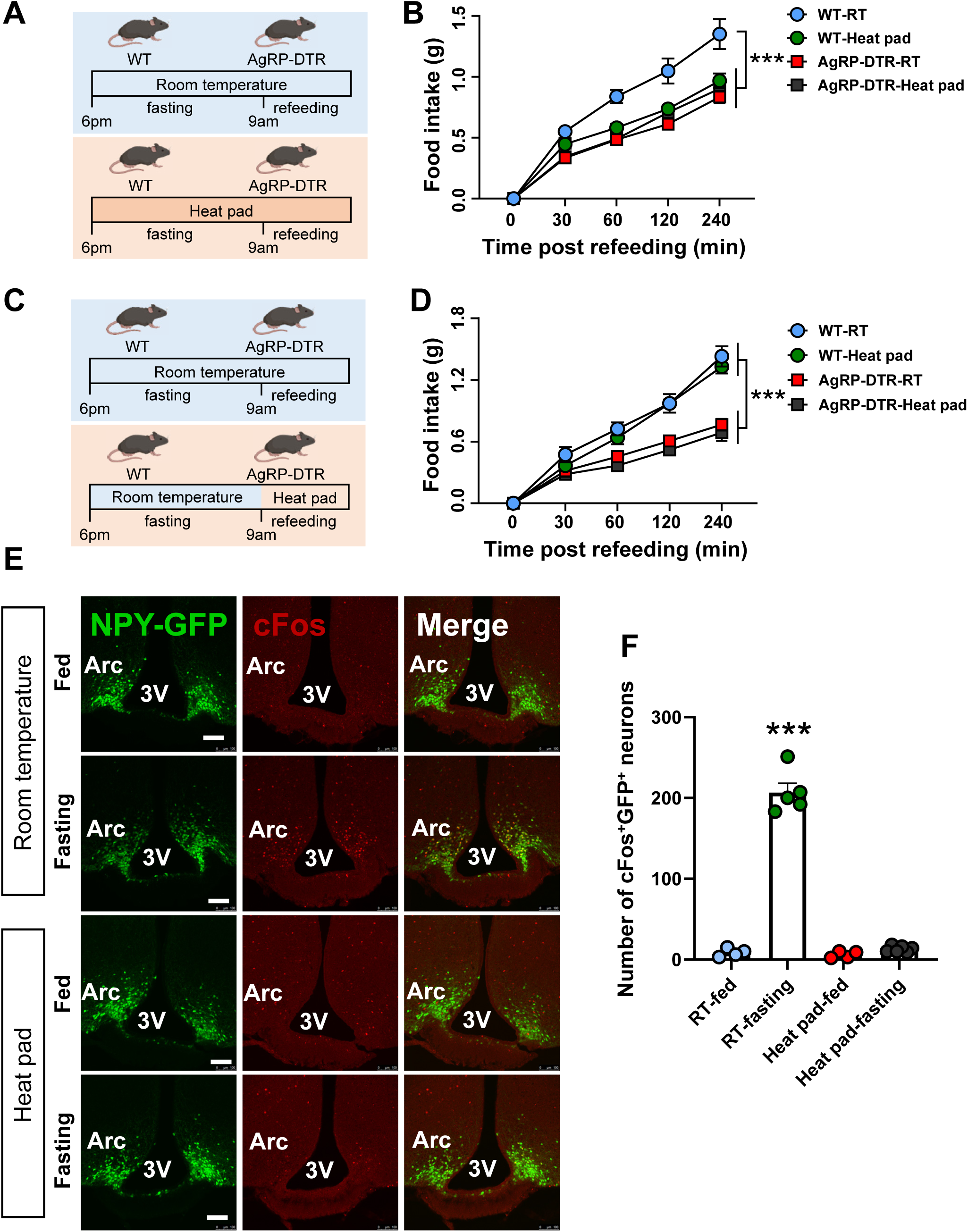
Contribution of AgRP neurons to body temperature maintenance during fasting. **(A, B)** WT and AgRP-DTR mice were fasted and refed in the room temperature or on a heat pad, as indicated in the schematic. (A) Schematic representation of the fasting–refeeding experimental paradigm. (B) Food intake post refeeding. Two-way repeated ANOVA followed by Bonferroni’s multiple comparisons test, ***p<0.001 between WT-RT and each of the other three groups. **(C, D)** WT and AgRP-DTR mice were fasted at room temperature, followed by refeeding either at room temperature or on a heating pad, as indicated in the schematic. (C) Schematic representation of the fasting–refeeding experimental paradigm. (D) Food intake post refeeding. Two-way repeated ANOVA followed by Bonferroni’s multiple comparisons test, ***p<0.001. n=6 for WT-RT/ WT-Heat pad/ AgRP-DTR-Heat pad groups. n=5 for AgRP-DTR-RT group in (B) and (D). **(E)** c-Fos immunostaining in the arcuate nucleus of NPY-GFP mice fasted overnight either at room temperature or on a heating pad. NPY neurons are shown in green, and cFos is shown in red. Scale bar = 100 μm. **(F)** Statistical comparison of cFos positive NPY neurons among different groups. Two-way repeated ANOVA followed by Bonferroni’s multiple comparisons test, ***p<0.001 between RT-fasting and RT-fed groups. n=4 for RT-fed group, n=5 for RT-fasting group, n=4 for Heat pad-fed group, n=6 for Heat pad-fasting group. All data represent the mean ± SEM.

AgRP neurons are known to be activated by fasting. To investigate the involvement of AgRP neuron activation in maintaining body temperature during fasting, we examined c-Fos activation in these neurons. As expected, in mice housed in room temperature, AgRP neurons exhibited increased c-Fos expression (Fig. 5E and 5F), which was completely eliminated when mice were housed with heating pad (Fig. 5E and 5F), suggesting that the main function of AgRP neuron activation during fasting is to promote energy intake for maintaining body temperature during fasting periods.

## Discussion

Homeotherms spend a considerable amount of energy to defend a much higher body temperature to the ambient room temperature for survival (Jensen *et al*., 2013; Reitman, 2018). This results in a conflicting situation during conditions with long-term food scarce and time-restricted feeding, in which there is a need to reduce energy expenditure to conserve energy reserve but also a need to spend necessary energy as heat production to defend body temperature (Cannon and Nedergaard, 2004). Although fasting is known to be linked with reduced energy expenditure (Jensen *et al*., 2013), the neural basis underlying body temperature maintenance during time-restricted feeding is not well understood. Our current results showed that lesions of AgRP neurons caused a rapid lethal phenotype during time-restricted feeding, suggesting that AgRP neurons are essential to mount adaptive behaviors to conditions with food scarce. One major defect is that these mice exhibited reduced food intake in response to food restriction. Thus, while control mice adaptively increased food intake during the period with food availability, mice with AgRP neuron lesion failed to do so. In support of this, AgRP neurons are also required to mount fasting-induced hyperphagia and to promote food anticipatory activity (Cai *et al*., 2023; Tan *et al*., 2014). This reduced feeding may be caused by a reduction in motivational feeding behaviors as the lesioned mice showed reduced food pellet retrieval by nose pokes in the FED3 cages. Strikingly, our results from the pair-fed feeding studies suggest that food intake in control mice, even at a much-reduced level matching that in the lesioned mice, is sufficient to maintain body temperature and survival. Thus, reduced food intake in the lesioned mice is not the cause of the observed lethality. Since the lethality was completely rescued by preventing body temperature drop with a heating pad, failure in body temperature maintenance is the cause of the lethality.

Our additional results from studies with acute fasting and cold exposure suggest that the lesioned mice, when provided food *ad libitum*, even with less food intake, showed completely normal body temperature maintenance, but failed to sustain body temperature maintenance when food was not available. In corroboration, in response to a warm environment with heating pad, while control mice reduced fasting-reeding, AgRP neuron lesion exhibited no difference in fasting refeeding between room temperature and heating pad, suggesting a role for AgRP neurons in promoting energy intake for body temperature maintenance. Supporting this, fasting-induced activation of AgRP neurons in room temperature was completely eliminated by switching to heating pad, reinforcing a physiological role for AgRP neuron activation in promoting energy intake for body temperature maintenance during fasting. These results collectively suggest a previously unknown role for AgRP neurons in defending body temperature maintenance during conditions with chronic food scarce including time-restricted feeding.

Of note, previous studies suggest that AgRP neurons, in addition to promoting feeding, have also been shown to reduce energy expenditure to promote positive energy balance (Cavalcanti-de-Albuquerque et al., 2019; Denis *et al*., 2015; Egawa et al., 1990; Ruan et al., 2014; Shi et al., 2017). However, these studies were mostly conducted in *ad libitum* feeding conditions with acute activation of AgRP neurons. Our results on AgRP neurons function in body temperature maintenance were observed under chronic food restriction or fasting with cold exposure, which is known to be associated with an increase in sympathetic nerve activity in thermogenic tissues coupled with increased activity of AgRP neurons (Brito et al., 2008). Our results on AgRP neurons in maintaining body temperature is supported by previous results showing that activation of these neurons promote energy expenditure (Qi *et al*., 2022), AgRP neurons are sensitive to changes in environmental temperature (Deem *et al*., 2020), and intermittent fasting rather increases energy expenditure (Liu et al., 2019). Thus, it appears that AgRP neurons serve as an integration site for both hunger and cold, leading to a coordinated regulation of feeding and thermogenesis for survival in response to challenging environments with food scarce and low ambient temperature.

While it is well accepted that activation of AgRP neurons is sufficient to promote feeding and body weight gain(Aponte *et al*., 2011; Krashes *et al*., 2011; Xu *et al*., 2018; Zhu *et al*., 2020), whether these neurons are required to maintain normal body weight regulation, especially under *ad libitum* HFD feeding conditions, highly relevant to human obesity therapeutics, remains controversial. While some animal models with various gene manipulations in AgRP neurons mimicking loss of function in these neurons exhibit reduced feeding or DIO (Cai *et al*., 2023; Miletta *et al*., 2020; Padilla *et al*., 2016; Tan *et al*., 2014), (Ajwani *et al*., 2024; Quiñones *et al*., 2018; Üner *et al*., 2015; Zhang *et al*., 2024), other similar animal models surprisingly exhibit obesity development(Denis *et al*., 2015; Joly-Amado *et al*., 2012; Qi *et al*., 2022). Our previous results from mice with AgRP neuron lesion suggest that these neurons are not required for normal body weight regulation when mice were fed *ad libitum* chow (Cai *et al*., 2023; Zhu *et al*., 2020), which is supported by recent results showing that no impact on the *ob/ob* obesity by complete disruption of neurotransmission from AgRP neurons (Cavalcanti de Albuquerque *et al*., 2025; Gou *et al*., 2025). Our current data showed that the same lesion caused no impact on DIO when HFD was provided *ad libitum* and produced no differences in obesity reduction in response to GLP1-R agonism.

The reasons underlying the discrepancies in DIO in these studies are not clear, but cannot be explained by potential developmental compensations as speculated by others (Bodur et al., 2025), since our lesion model is adult onset. As AgRP is also expressed in a few peripheral sites (Leon-Mercado et al., 2025; Liu et al., 2021; Liu et al., 2023), it is conceivable that the differences in DIO from those models with gene manipulations/deletion mediated by AgRP-Cre mouse lines may be contributed from those peripheral sites. In addition, given the demonstrated role of AgRP neurons in sensing ambient temperature and controlling body temperature maintenance, it is also conceivable that the metabolism in mice with comprised function of AgRP neurons may show altered responses to variations in housing conditions including temperature and accessibility of food, leading to the observed slightly reduced (Cai *et al*., 2023; Miletta *et al*., 2020; Padilla *et al*., 2016; Tan *et al*., 2014), (Ajwani *et al*., 2024; Quiñones *et al*., 2018; Üner *et al*., 2015; Zhang *et al*., 2024) or increased DIO (Denis *et al*., 2015; Gupta et al., 2017; Qi *et al*., 2022), and this phenomenon has been reported previously (Reimúndez et al., 2018). Given our model with almost complete lesion of AgRP neurons specifically in the brain, our results strongly support that AgRP neurons are not required for body weight regulation in *ad libitum* feeding conditions and argue that AgRP neurons may not be used as an effective anti-obesity target.

In sum, our current results demonstrate that AgRP neurons are essential for survival during time-restricted feeding but not required for normal body weight regulation when mice fed *ad libitum* with either chow or HFD food. This concept is supported by earlier findings that the melanocortin pathway is biased toward one-directional toward promoting positive energy balance (Li et al., 2023), and is also consistent with the idea that leptin functions rather as a fasting hormone, and the reduced levels of leptin during fasting conditions leads to activation of AgRP neurons for the adaptation to food scarce environments (Ahima et al., 1996). The involvement of these neurons in adaptive behaviors to chronic food deficient states is also supported by previous results involving these neurons in behavioral changes in anorexia models (Kucukdereli et al., 2024; Miletta *et al*., 2020; Sutton Hickey et al., 2023).

Here we used the animal model with AgRP neuron lesion and provide a novel insight on the role of AgRP neurons in promoting body temperature maintenance during chronic food restriction. However, this model, although provides a powerful tool to examine the necessity of AgRP neurons in fasting, cannot be used to study how AgRP neurons are engaged in the control of body temperature during the progression of feeding restriction. Further studies with monitoring and altering AgRP neuron activity will reveal physiological relevance of AgRP neurons in body temperature maintenance in response to feeding restriction or fasting with cold.

## Materials and Methods

### Animals

Mice were housed in the Institute of Molecular Medicine animal facility. They were maintained at 21-22 °C on a 12 h light/12h dark cycle with standard pellet chow (Teklad F6 Rodent Diet 8664, 4.05 kcal/g, 12.5% kcal from fat, Harlan Teklad) and water ad libitum-except in fasting experiments and high fat diet challenging experiments. Animal care and procedures were approved by University of Texas Health Science Center at Houston Institutional Animal Care and Use Committee. NPY-GFP (JAX: 006417) were purchased from the Jackson Laboratory. AgRP-DTR mice were provided by Q. Wu of Baylor College of Medicine. NPY-GFP and AgRP-DTR mice were crossed and bred together to allow GFP expression in AgRP/NPY neurons in the ARC. All experiments were performed on adult mice (8–10 weeks old, both male and female). Mice were euthanized when body weight dropped by >20%, as mandated by the ethics committee.

### Intracerebroventricular injections

Intracerebroventricular (i.c.v) injection was conducted to deliver the DTX (#D0567, Sigma-Aldrich) to the cerebrospinal fluid (CSF) in the lateral ventricles. Briefly, mice were anesthetized with a ketamine/xylazine cocktail (100 mg/kg and 10 mg/kg, respectively), and their heads were fixed to a stereotaxic apparatus. 0.1 µL saline or DTX solution (50 ng/ µL) were delivered through a 0.5 µL syringe (Neuros Model 7000.5 KH, point style 3; Hamilton, Reno, NV, USA) mounted on a motorized stereotaxic injector (Quintessential Stereotaxic Injector; Stoelting, Wood Dale, IL, USA) at a rate of 20 nL/min. The coordinates to target the right lateral ventricle was: anteroposterior (AP): 0.0 mm; mediolateral (ML): +1.0 mm; dorsoventral (DV): -2.5 mm. All i.c.v DTX injections were unilateral since CSF can circulate through the whole brain.

### Body weight measurement and semaglutide treatment

Mice were put on HFD (Research Diets D12492, 20% protein, 60% fat, 20% carbohydrate, 5.21 kcal/g) three weeks after i.c.v injection. Long-term body weights were measured weekly for 8 weeks. Then, mice were received twice subcutaneous (s.c) injection of 40 µg/kg semaglutide (#29969, Cayman) for one week and daily for the next two weeks. Body weights were measured daily during semaglutide treatment.

### Metabolic cages

Mice were individually housed in chambers of the PhenoMaster cages (TSE systems, Chesterfield, Missouri, USA). Mice were given ad libitum access to a normal chow diet or HFD and water. Food intake, O2 consumption and locomotion activity levels were measured using indirect calorimetry continuously at different time points for one week. Data was averaged through different time points in light and dark cycles, respectively, for comparison. The data from the first two days and the last day was removed.

### Body temperature measurement

Mice were anesthetized with isoflurane and then subcutaneously implanted with a transponder (TP-1000, BMDS) on the back above the brown adipose tissue. Body temperature was measured using a pocket reader (DAS-8027-IUS, BMDS).

### Food restriction

Four weeks after a single i.c.v injection of DTX, WT and AgRP-DTR male mice were subjected to a food restriction, whereby the mice were granted access to chow only during 6 pm to 11 pm. Food intake, body weight, body temperature and survival were recorded daily during the experiment.

### Heating pad treatment

In the present study, mice were placed on a heating pad set to 36 °C to maintain the cage temperature at approximately 30 °C, which is within the thermoneutral zone for mice.

### Feeding Experimentation Device 3 (FED3) experiment

Four weeks after a single i.c.v injection of DTX, WT and AgRP-DTR male mice were singly housed for one week. Mice were then given access to food pellets delivered by the FED3 system (Open Ephys, https://hackaday.io/project/106885-feeding-experimentation-device-3-fed3) in their home cages. Each FED3 device was loaded with 20 mg dustless precision pellets (F0163, Bio-Serv). During the first week, the operant schedule was set to a fixed ratio of 1 (FR=1, one nose poke per pellet), and in the second week, it was increased to a fixed ratio of 5 (FR=5, five nose poke per pellet). Food intake and body weight were monitored daily.

### Pair feeding

Pair feeding was performed by measuring the daily food intake of the group with lower consumption and providing the same amount of food to the other group. Mice were singly housed for one week before the initiation of the pair-feeding experiment. During the experiment, food was available only between 6 pm and 11 pm. Mice in the AgRP-DTR–Heat pad and WT–RT groups were provided with the same amount of food consumed by the AgRP-DTR–RT group. Food intake, body weight, body temperature, and survival were recorded daily.

### Cold exposure experiment

Mice were singly housed for one week prior to the experiment. Water was available ad libitum, but bedding was removed during cold exposure. Mice were exposed to 4 °C from 6 pm to 11 pm with access to food. One week later, the same mice were exposed to 4 °C from 6 pm to 11 pm without food. Food intake and body temperature were monitored hourly.

### Fasting-refeeding test

WT and AgRP-DTR mice were fasted overnight and subsequently refed under the same temperature conditions (room temperature or heating pad set to 36 °C; Fig. 5A). One week later, the same mice were fasted at room temperature and then refed either at room temperature or on a heating pad (Fig. 5C). Food intake was measured at 30, 60, 120, and 240 minutes after refeeding.

### Immunostaining and imaging

Mice were anesthetized with a ketamine/xylazine cocktail (100 and 10 mg/kg, respectively) and transcardially perfused with 0.9% normal saline followed by 10% formalin. Freshly fixed brains were collected and placed in 10% formalin at room temperature overnight for post-fixation, then dehydrated in 30% sucrose solution. Brains were frozen and sectioned onto 30 μm thickness slices with a sliding microtome. Brain sections were immunostained with rabbit anti-cFos (1:1000, 2250 S, Cell Signaling Technology) overnight. Then, all brain sections were incubated with Alexa Fluor 568 donkey anti-rabbit IgG (1:500, A10042, Thermo Fisher Scientific). Brain sections with NPY-GFP and red fluorescence were visualized with confocal microscopy (Leica TCS SP5, Leica Microsystems, Wetzlar, Germany). cFos were counted from four matched sections containing rostral, middle, and caudal of arcuate from individual mice.

## Statistical Analysis

All the data were recorded and organized using Excel sheets first and then exported to GraphPad Prism (GraphPad Software, Inc., La Jolla, CA, USA). Two-way ANOVA followed by Bonferroni’s multiple comparisons were used. For single variable comparisons, such as food intake measurements, unpaired two-tailed Student’s t tests were used. Error bars in all graphs are presented as mean±SEM.

## Acknowledgements

We acknowledge Dr. Qi Wu for providing the AgRP-DTR mice. Q.T. is supported by NIH R01 DK136284, R01 DK 135212 and R01 DK 131466 (QT), and R01DK109934 and DOD HT94252310156 (QT and BRA). The project is benefited from the Optogenetics and Viral Vectors Core, supported by NIH IDDRC grant 1 U54 HD083092, and the Gene Vector Core, at Baylor College of Medicine for providing viral preparations. QT is the holder of the Cullen Chair in Molecular Medicine and Hans J. Eberhard MD, PhD and Irma Gigli, MD Distinguished Chair in Immunology at McGovern Medical School.

## Author contribution

The experiments were mainly conducted by C.S. with helps from Y.X. B.R.A. provided essential reagents. C.S. wrote the manuscript with significant inputs from B.R.A., and Q. T.

## Supplementary Materials

Figs. S1 to S2

