## Supplementary Figures for "An obligatory role for AgRP neurons in maintaining body temperature during time-restricted feeding"

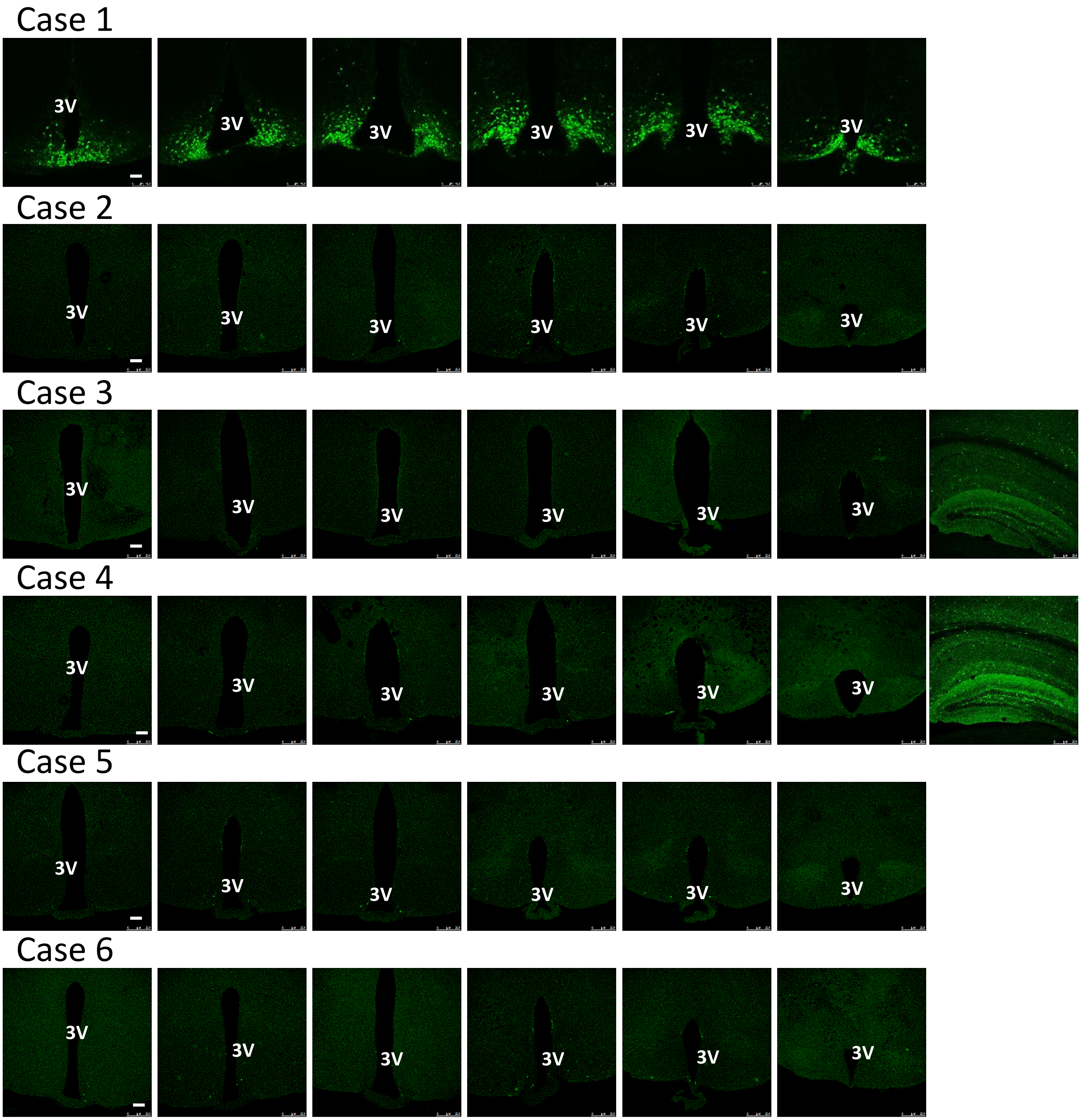


**Supplementary Fig.1. Representative cases of mouse brains with AgRP neuron lesion.** Case 1 is the control of NPY-GFP expression and cases 2-6 are NPY-GFP::AgRP-DTR mice received 5ng DTX icv. Note that a few scattered neurons remain with GFP expression, which may represent those AgRP-negative NPY neurons. In Cases 3 and 4, many GFP NPY positive neurons in the hippocampus were shown. The sections in all cases were presented from a rostral to caudal order. Scale bar= 50uM. 3V: the third ventricle.

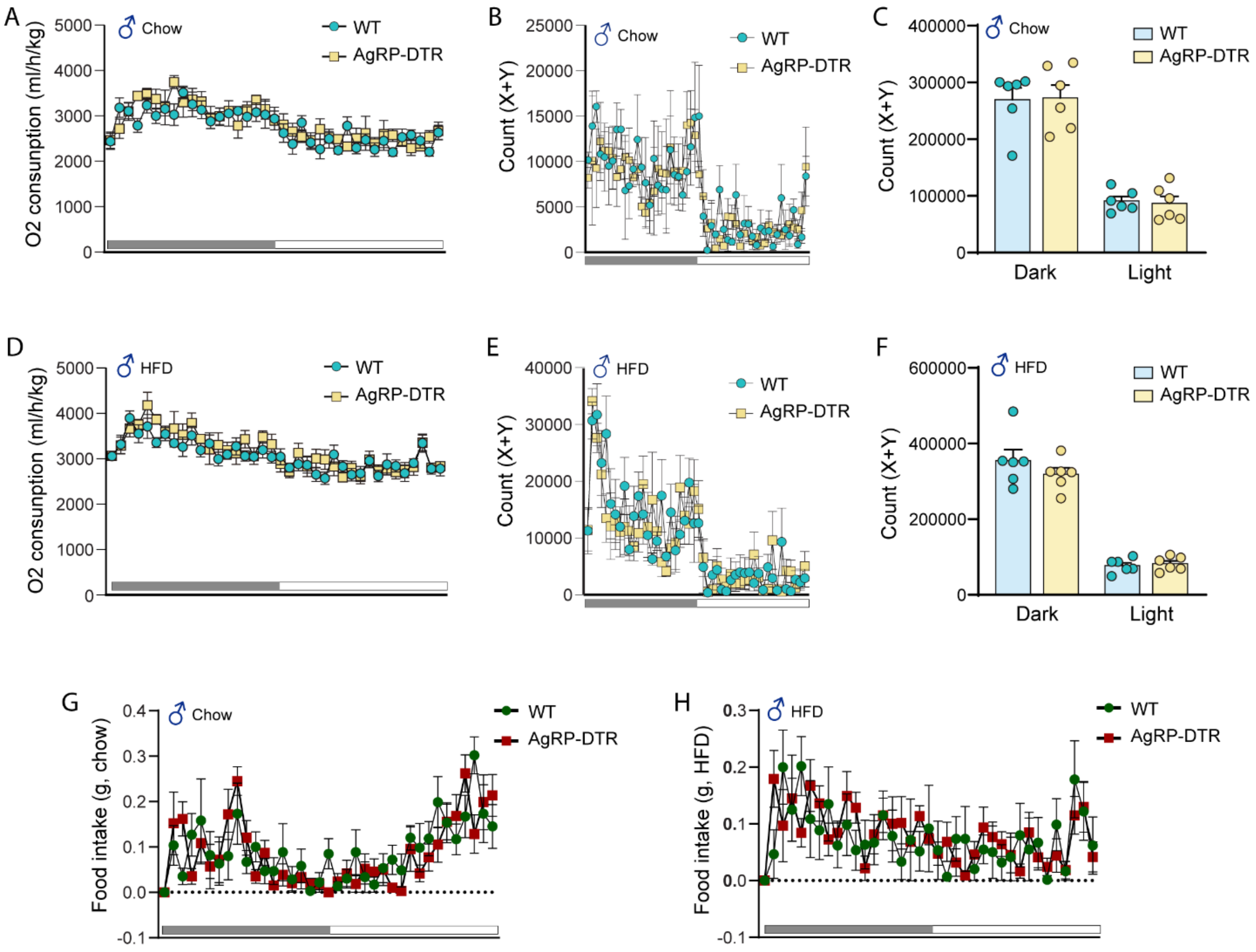


**Supplementary Fig.2 AgRP neurons are not required in normal energy balance responses to chow and HFD feeding.**

Comparisons in O2 consumption after ablation of AgRP neurons on chow (A) and HFD (D). Real-time locomotor activity patterns of male mice measured by the TSE metabolic cages on chow (B) and HFD (E). Comparisons in the locomotor activity levels of mice fed on chow (C) and HFD (F) during the day and night periods. Real-time food intake patterns of mice on chow (G) and HFD (H). All data is presented as mean±SEM, n=6 mice per group.
